# *Drosophila fabp* is a retinoid-inducible gene required for Rhodopsin-1 homeostasis and photoreceptor survival

**DOI:** 10.1101/2021.04.16.440122

**Authors:** Huai-Wei Huang, Hyung Don Ryoo

## Abstract

Retinoids act as chromophore co-factors for light-detecting rhodopsin proteins. In vertebrates, retinoids also actively regulate gene expression. Whether retinoids regulate gene expression in *Drosophila* for a specific biological function remains unclear. Here, we report that *Drosophila fatty acid binding protein* (*fabp*) is a retinoid-inducible gene required for Rhodopsin-1 (Rh1) protein homeostasis and photoreceptor survival. Specifically, we performed a photoreceptor-specific gene expression profiling study in flies bearing a misfolding-prone Rhodopsin-1 (Rh1) mutant, *ninaE*^*G69D*^, which serves as a *Drosophila* model for Retinitis Pigmentosa. *ninaE*^*G69D*^ photoreceptors showed increased expression of genes that control Rh1 protein levels, along with a poorly characterized gene, *fabp*. We found that in vivo *fabp* expression was reduced when the retinoids were deprived through independent methods. Conversely, *fabp* mRNA was induced when we challenged cultured *Drosophila* cells with retinoic acid. In flies reared under light, loss of *fabp* caused an accumulation of Rh1 proteins in cytoplasmic vesicles. *fabp* mutants exhibited light-dependent retinal degeneration, a phenotype also found in other mutants that block light-activated Rh1 degradation. These observations indicate that a retinoid-inducible gene expression program regulates *fabp* that is required for Rh1 proteostasis and photoreceptor survival.

**Author Summary:** Rhodopsins are light-detecting proteins that use retinoids as chromophore co-factors. In vertebrates, retinoids also actively regulate gene expression. Whether retinoids regulate Rhodopsin function aside from its role as a chromophore remains unclear. Here, we report that *Drosophila fatty acid binding protein* (*fabp*) is a retinoid-inducible gene required for Rhodopsin-1 (Rh1) protein homeostasis and photoreceptor survival. Specifically, we found that *fabp* is among the genes induced by a misfolding-prone Rhodopsin-1 (Rh1) mutant, *ninaE*^*G69D*^, which serves as a *Drosophila* model for Retinitis Pigmentosa. We further found that *fabp* induction in *ninaE*^*G69D*^ photoreceptors required retinoids. *fabp* was required in photoreceptors to help degrade light-activated Rh1. In the absence of *fabp*, Rh1 accumulated in cytoplasmic vesicles in a light-dependent manner, and exhibited light-dependent retinal degeneration. These observations indicate that a retinoid-inducible gene expression program regulates *fabp* that is required for Rh1 proteostasis and photoreceptor survival.

## Introduction

Rhodopsins are G-protein coupled proteins associated with retinal chromophores to detect light and initiate signal transduction (1). As in mammals, *Drosophila* has multiple Rhodopsins, including *ninaE* (*neither inactivation nor afterpotential*) that encodes the Rhodopsin-1 (Rh1) protein expressed in R1 to R6 photoreceptors (2-4). Functional Rh1 is covalently attached to the 11-cis-3-hydroxyretinal chromophore, which is derived from dietary vitamin A (5-7). *ninaE* loss of function results in an impairment of light detection (4, 8).

Abnormal Rh1 protein homeostasis is a frequent cause of retinal degeneration. One class is caused by a group of *ninaE* missense mutations that dominantly cause progressive age-related retinal degeneration (9, 10). These alleles are analogous to human rhodopsin mutations that underlie age-related retinal degeneration in Autosomal Dominant Retinitis Pigmentosa (ADRP) patients (11, 12). Using the *Drosophila ninaE*^*G69D*^ allele as a model, we previously established that these mutations impose stress in the endoplasmic reticulum (ER), which contributes to retinal degeneration (13, 14). The human rhodopsin allele that is most frequently found associated with ADRP, the P23H mutant, similarly causes ER stress in mammalian cells (15).

Cellular mechanisms that regulate rhodopsin protein levels affect retinal degeneration. Flies bearing one copy of the *ninaE*^*G69D*^ allele have total Rh1 protein levels reduced by more than half, indicating that both the mutant and the wild type Rhodopsin-1 proteins undergo degradation in these flies (9, 10). Three ubiquitin ligases that specialize in the degradation of misfolded endoplasmic reticulum (ER) proteins mediate the degradation of Rh1 in *ninaE*^*G69D*^ flies (16). Overexpression of these ubiquitin ligases can delay the onset of retinal degeneration in *Drosophila ninaE*^*G69D*^ flies, indicating that excessive misfolded Rh1 is a contributing factor to retinal degeneration (14, 16).

Functional wild type Rh1 proteins also undergo degradation after being activated by light. Specifically, this occurs after light-activated Rh1, also referred to as metarhodopsin (M), engages with Arrestin that mediates feedback inhibition (17). Rh1 forms a stable complex with Arrestin and together undergo endocytosis for degradation (18-21). Such degradation of activated Rh1 is essential, as too much Rh1 accumulation in the endosome/lysosome defective photoreceptors results in light-dependent retinal degeneration (18, 22-26). In Rh1 endocytosis/degradation defective mutants, retinal degeneration could be delayed by conditions that reduce overall Rh1 levels (24, 25, 27), indicating that too much active Rh1 is a cause of retinal degeneration. These aspects appear to be conserved across phyla, as the human rhodopsin mutants that exhibit high affinities for Arrestin display endosomal abnormalities and are associated with severe forms of ADRP (28, 29).

Retinoids are among the molecules implicated in regulating Rh1 protein levels. Deprivation of vitamin A, which serves as a precursor for the retinal chromophore, causes a reduction in overall Rh1 levels (30-34). Such an effect is largely attributed to the importance of chromophores in Rh1 protein maturation. Aside from its role as a rhodopsin cofactor, retinoids regulate gene expression in vertebrates. The best characterized transcription factors that mediate this response are nuclear hormone receptors, including RAR and RXR, which become transcriptional activators upon binding to all-trans or 9-cis retinoic acid (35). Cellular Retinoic Acid Binding Protein -I and -II (CRABP-1, -II) bind to the lipophilic retinoic acids and deliver them to RAR and RXR in the nucleus (36, 37). In addition to mediating the RA signaling response, CRABP-II itself is induced by RA signaling (38). Whether these mechanisms are conserved in *Drosophila* has not been examined in detail in part because the *Drosophila* genome does not encode a RAR homolog (7, 39). Intriguingly, several *Drosophila* genes require the retinoid precursor vitamin A for their gene expression (40, 41). One such gene is *highroad*, whose expression is induced by retinoic acids in cultured cells and mediates the degradation Rh1 in *ninaE*^*G69D*^*/+* (42). These results suggested that retinoid-mediated gene expression programs may be involved in Rh1 homeostasis.

Here, we report that Rh1 protein levels are regulated by the *Drosophila* CRABP homolog, *fatty acid binding protein* (*fabp*). Similar to CRABP-II, *fabp* expression is induced by retinoids. Loss of *fabp* enhances total Rh1 levels in *ninaE wild type* and *G69D* mutant backgrounds. Moreover, loss of *fabp* causes light-dependent retinal degeneration. Our results indicate that this retinoic acid inducible gene controls Rh1 protein levels, and this regulatory axis is essential for photoreceptor survival.

## Results

### Photoreceptor-specific gene expression profiling shows *fabp* induction in *ninaE*^*G69D*^ eyes

To better understand how photoreceptors respond to stress imposed by the *ninaE*^*G69D*^ allele, we performed a photoreceptor-specific gene expression profiling analysis. We specifically employed a previously described approach in which the expression of the nuclear envelope-localized *EGFP::Msp300*^*KASH*^ is driven in specific cell-types through the Gal4/UAS system to isolate the EGFP-labeled nuclei for RNA-seq analysis (43, 44). We used the *Rh1-Gal4* driver to isolate *ninaE* expressing R1 to R6 photoreceptor nuclei from the adult fly ommatidia (Figure 1A). Microscopy imaging confirmed that anti-EGFP beads enriched the EGFP::Msp300^KASH^-tagged nuclei (Figures 1B, C). RNA-seq was performed with nuclei isolated from *ninaE wild type* and *ninaE*^*G69D*^*/+* photoreceptors.

**Figure 1:**
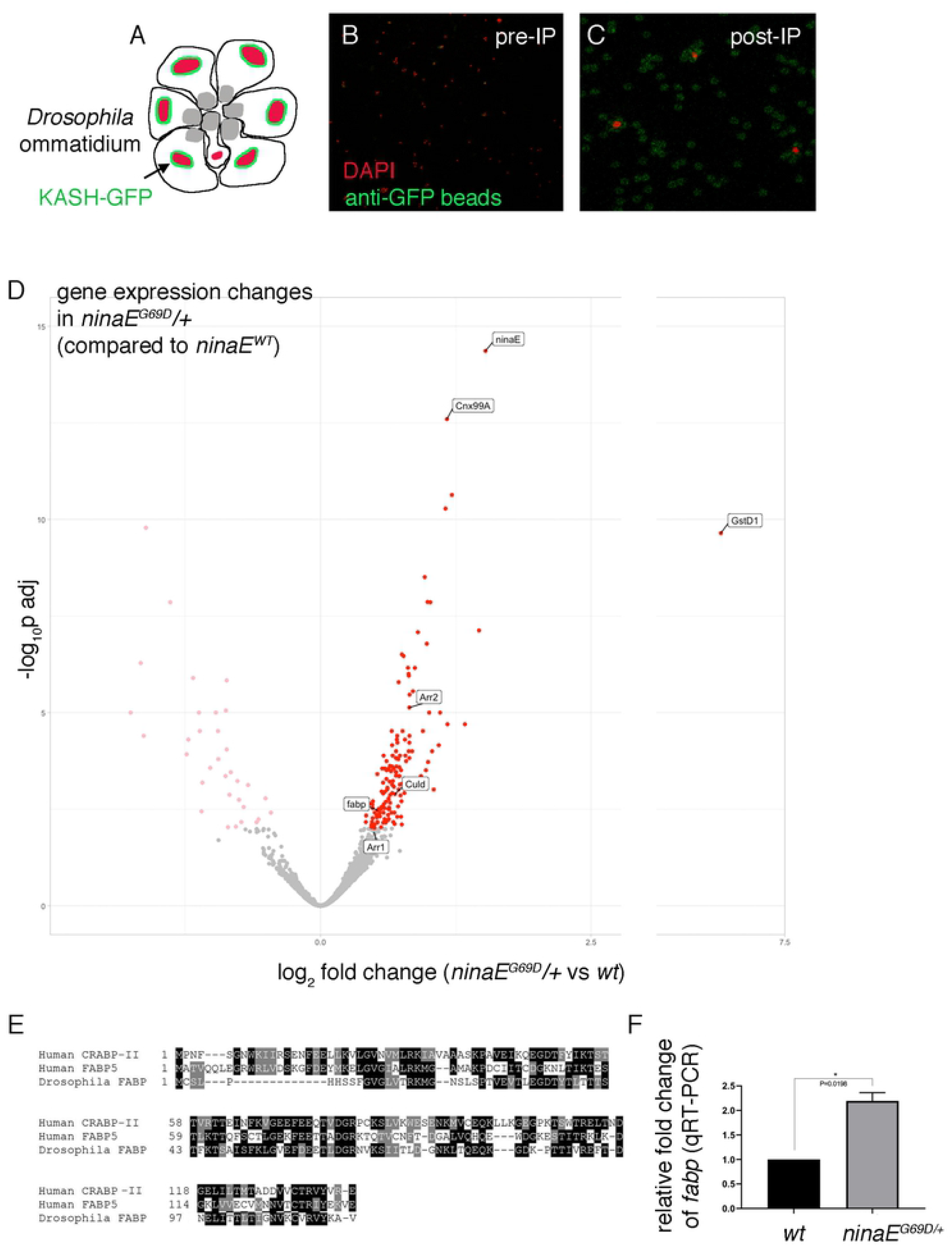
Photoreceptor-specific gene expression profiling in *ninaE*^*G69D*^ eyes. (A) A schematic diagram of the *Drosophila* ommatidium. Shown are seven photoreceptor cells, R1 to R7. KASH-GFP (green) coats the outer membranes of R1 to R6 nuclei (red). Rh1 localizes to the apical membrane structure known as rhabdomeres (gray). (B, C) Purification of photoreceptor nuclei tagged with EGFP-Msp300^KASH^ (KASH-GFP). *Rh1-Gal4* was used to express *uas-KASH-GFP* in R1 to R6 photoreceptors. Nuclei are labeled with DAPI (red) and the anti-GFP beads are in green. (B) Before anti-GFP bead purification. (C) After purification, the DAPI labeled nuclei are associated with the beads. (D) Volcano plot of differential gene expression compared between *ninaE wild type* and *ninaE*^*G69D*^*/+* photoreceptors. The y axis shows -log_10_ (adjusted p value). The x axis represents log_2_ fold change, with those whose expressions increase in *ninaE*^*G69D*^*/+* on the right (adjusted p<0.01 are labeled in red). Log_2_ fold change above 2.5 is not in scale. Genes with nonsignificant changes (adjusted p>0.01) are in gray. (E) Sequence comparison between *Drosophila* FABP, human CRABP-II and FABP5. (F) qRT-PCR results of *fabp* from *ninaE wild type* (left) and *ninaE*^*G69D*^*/+* fly heads. Error bars represent Standard Error (SE). t-test was used to assess statistical significance. * = p<0.05.

Differential gene expression analysis showed 182 genes whose expression changed with adjusted p values below 0.01 (Supplementary Table 1). Among the most highly induced genes was *gstD1* (Figure 1D), which was also identified as an ER stress-inducible gene in a separate study performed with larval imaginal discs (Brown et al., under review). The ER chaperone encoding *cnx99A* was also induced (Figure 1D), consistent with the previous report that *ninaE*^*G69D*^ imposes ER stress in photoreceptors (13).

Also, notable from the differential gene expression analysis was the induction of genes that could affect Rh1 levels. *ninaE* was itself induced in *ninaE*^*G69D*^ samples (Figure 1D). Since *ninaE*^*G69D*^*/+* flies have very low Rh1 levels (9, 10), we speculate that increases in *ninaE* transcription may be part of a feedback homeostatic response. Also induced were genes *Arrestin 1* (*Arr1*), *Arrestin 2* (*Arr2*) and *culd* (Figure 1D), which promote the degradation of light-activated Rh1 in photoreceptors (17, 18, 23, 45).

Also, among the *ninaE*^*G69D*^-induced genes was *fabp* (Figure 1D), which encodes a protein homologous to human CRABP-1, -II and FABP5 (Figure 1E). We validated the induction of *fabp* mRNA in *ninaE*^*G69D*^*/+* through q-RT PCR (Figure 1F). The human homologs of *fabp* are known to bind all trans RA with high affinity (36, 37, 46, 47). Notably, CRABP-II is one of the well-characterized RA inducible genes in mammalian cells (38). *fabp* drew our interest because a retinoic acid-inducible *Drosophila* gene, *hiro*, regulates Rh1 levels in *ninaE*^*G69D*^*/+* flies (42).

### *fabp* expression is regulated by Vitamin A and retinoids

To test if *Drosophila fabp* is also regulated by retinoic acids (RA), we examined *fabp* mRNA levels through RT-qPCR in *Drosophila* S2 culture cells treated with or without 10mM RA. We found that RA treated cells showed an increase in *fabp* transcripts after 60 minutes of RA exposure (Figure 2A, B).

**Figure 2:**
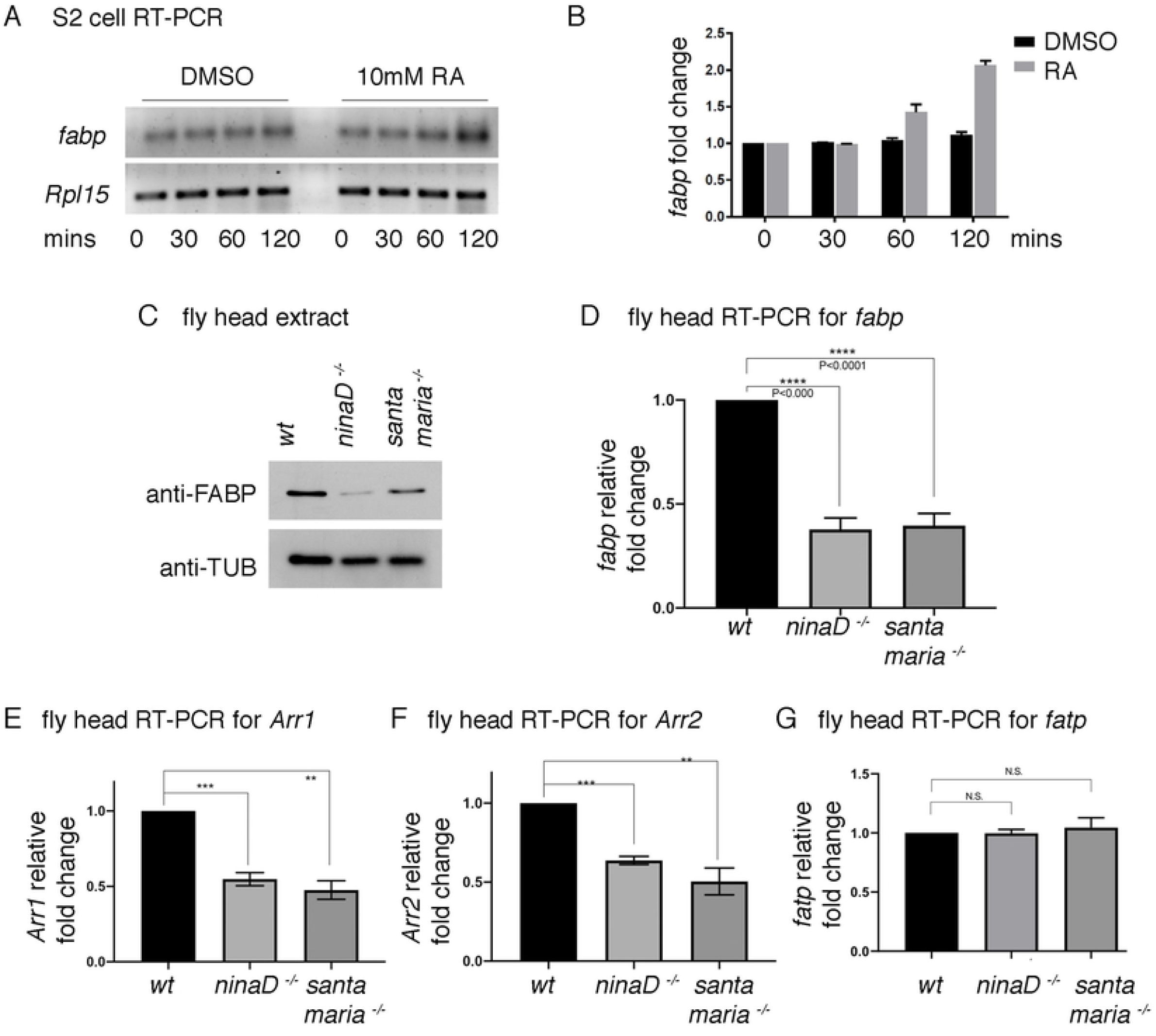
*fabp* expression is regulated by carotenoids and retinoic acid. (A) The time course of *fabp* mRNA induction as assessed through semi-quantitative RT-PCR for *fabp* (top gel) and the control *Rpl15* (bottom gel). Cultured *Drosophila* S2 cells were treated with either control DMSO (left 4 lanes) or with 10mM all-trans retinoic acid (RA, right 4 lanes) for the indicated periods of time. (B) qRT-PCR of *fabp* from S2 cells treated with DMSO (black) or 10mM all-trans retinoic acid (grey bars) for the indicated period of time. The y axis shows fold induction as compared to the results from the DMSO controls. (C) Western blot of FABP (top gel) and β-Tubulin (bottom gel) from adult fly head extracts of the indicated genotypes. *w*^*1118*^flies were used as *wild type* controls. (D-G) qRT-PCR-based assessment of indicated mRNAs from adult fly heads of the indicated genotypes. The levels of *fabp* (D), *Arr1* (E), *Arr2* (F), *fatp* (G) are shown. The y axis shows fold changes compared to results obtained from *wild type* control samples. In all qRT-PCRs, *RpL15* qRT-PCR results were used to normalize the levels of transcripts of interest. Error bars represent standard error (SE). Statistical significance was 599 assessed through two tailed t-tests. ** = p<0.005, *** = p<0.0005, **** = p<0.0001.

To examine if *fabp* expression in fly tissues is affected by the availability of Vitamin A and its metabolites, we examined *fabp* levels in the mutants of *ninaD* and *santa maria* that have impaired transport of carotenoids, the precursors of retinoids. These mutants are devoid of retinoids in the retina, as evidenced by defective rhodopsin maturation and light detection (31, 34). We found that these mutants had reduced FABP protein as assessed through western blot of fly head extracts (Figure 2C). Consistently, the mutants also had reduced *fabp* mRNA levels as assessed through q-RT-PCR (Figure 2D).

These observations prompted us to examine if other *ninaE*^*G69D*^-inducible genes require *santa maria* and *ninaD* for proper expression. We focused on candidates known to be involved in Rh1 protein regulation. Among those tested, the mRNAs of *Arr1* and *Arr2* were found to be reduced in *ninaD* or *santa maria* mutant backgrounds (Figure 2E, F). Not all genes involved in Rh1 homeostasis were affected in these mutants. For example, *fatty acid transport protein* (*fatp*) is a gene whose loss-of-function increases Rh1 protein levels (27). *fatp* mRNA levels were affected neither in the mutant backgrounds of *ninaD* nor *santa maria* (Figure 2G). These results indicate that the expression of *fabp, Arr1, and Arr2* specifically require retinoid and carotenoid transporters, *ninaD* and *santa maria*.

### An *fabp* protein trap line shows carotenoid-dependent expression in the larval intestine

To independently validate the carotenoid-dependent expression of *fabp in vivo*, we utilized the *fabp* protein trap line *CA06960*. This P-element insertion line has a GFP with splice donor and acceptor sites, designed to make fusion proteins with the endogenous *fabp* coding sequence (Figure 3A). Anti-GFP western blot of fly extracts confirmed the expression of a GFP-fused protein in adult fly head extracts with the predicted size (Figure 3B).

**Figure 3:**
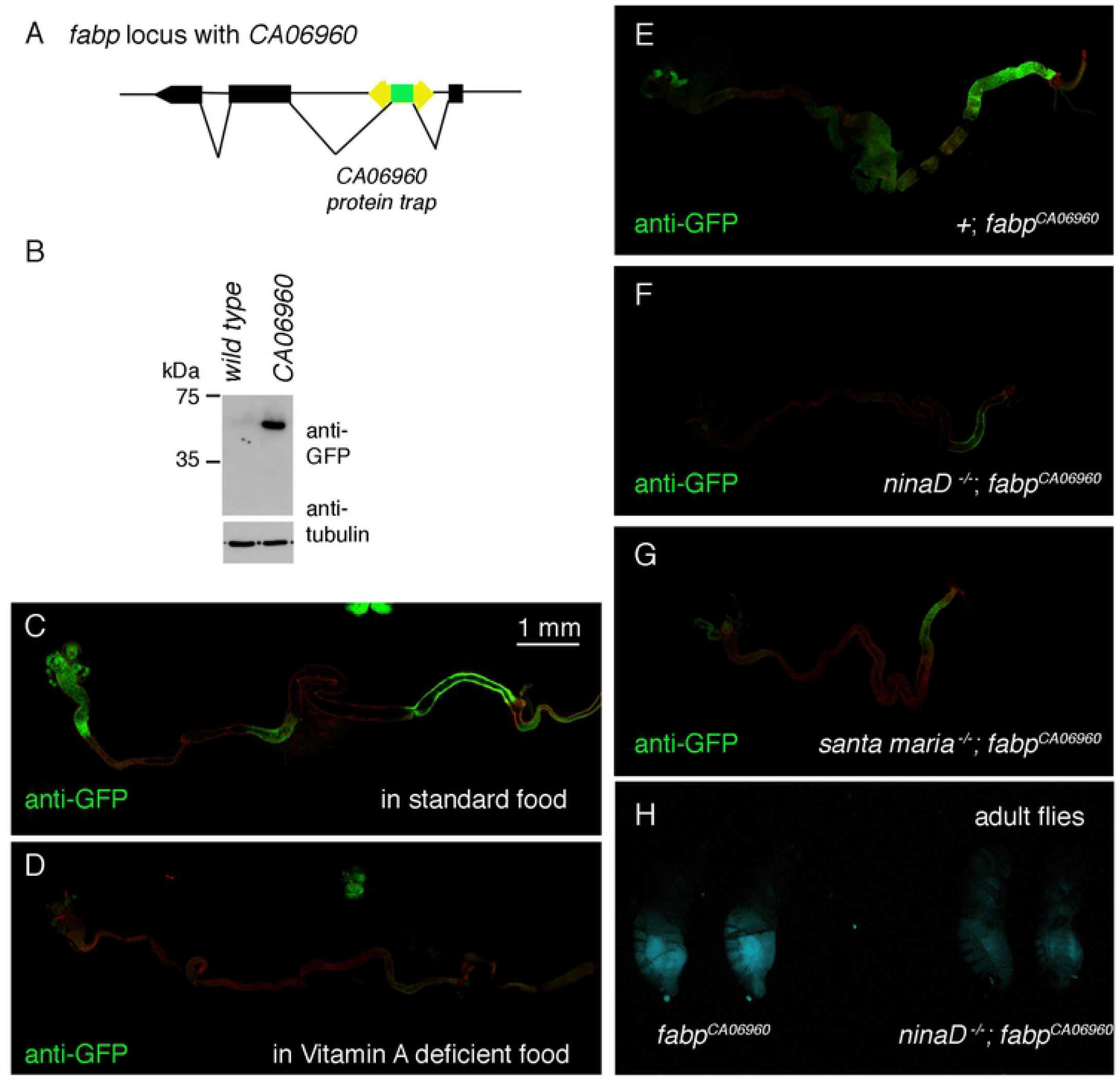
The expression of an *fabp* GFP protein trap line requires Vitamin A and its transporter genes. (A) The structure of the *fabp* locus and the *CA06960* protein trap line. (B) Anti-GFP western of adult head extracts from control and the *CA06960* protein trap line. (C-G) Images of dissected *fabp*^*CA06960*^ third instar larval intestines immuno-labeled with anti-GFP antibody (green). GFP signal is detected in distinct regions of the intestine in flies reared under standard conditions (C), which decreases in those reared in a Vitamin A deficient food (D). (E-G) GFP signal from flies reared under standard food in the control genotype (E), in the *ninaD*^*1*^ mutants (F) and in the *santa maria*^*1*^ mutant background (G). (H) GFP signal of *fabp*^*CA06960*^ adult females in the control genetic background (left two flies) and in the *ninaD*^*1*^ mutant background (right two flies). The scale bar in C is for images C-G.

In the third instar larva, the *fabp*^*CA06960*^ line had GFP expression detectable in several regions of the intestine (Figure 3C, E). Such expression was abolished when the flies were reared in Vitamin A deficient food (Figure 3D). Consistently, the expression of GFP was suppressed in the mutant backgrounds of *ninaD* and *santa maria* (Figure 3F, G). In adult flies, the GFP signal was most prominent in the female abdomen, which was reduced in the *ninaD* mutant background (Figure 3H). These results independently support the idea that *fabp* expression depends on carotenoids.

### Loss of *fabp* increases Rh1 protein levels

To test possible Rh1 regulation by retinoids, we examined several *Drosophila* homologs of mammalian RA signaling mediators. The candidate genes we analyzed included enzymes that convert Vitamin A to retinoids (e.g. *ninaB* (48)), nuclear hormone receptors (e.g. knl and eg) and *fabp*. For our assay, we used *ninaE*^*G69D*^*/+* flies, which have drastically reduced Rh1 protein levels as compared to *ninaE wild type* flies (Figure 4A). Specifically, we drove the expression of RNAi lines that target the genes of interest in the photoreceptors of these *ninaE*^*G69D*^*/+* flies using the Rh1-Gal4/UAS system (Figure 4A). An RNAi line that targeted *fabp* showed a reproducible effect of partially enhancing Rh1 levels as assessed through western blots of fly head extracts (Figure 4A, B).

**Figure 4:**
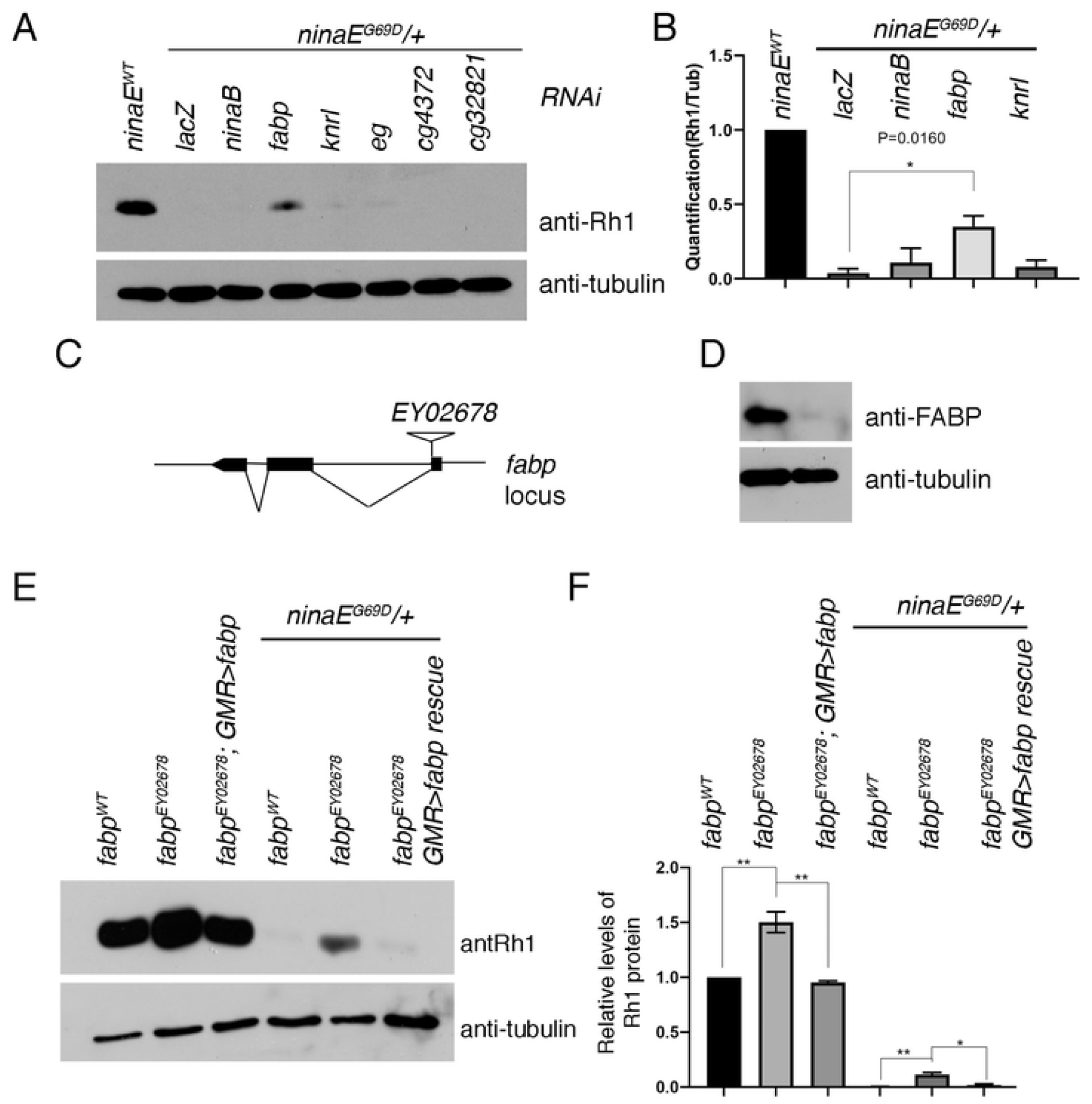
loss of *fabp* enhances Rhodopsin-1 protein levels. (A, B) Western blot of Rhodopsin-1 (Rh1) and β-tubulin from fly heads extracts. (A) The first lane is from *ninaE wild type* samples. The remaining lanes are from *ninaE*^*G69D*^*/+* flies with the indicated genes knocked down through RNAi with the Rh1-Gal4 driver. *lacZ* RNAi (lane 2) was used as a negative control. (B) Quantification of relative Rh1 band intensities as normalized to β-tubulin. (C) A schematic diagram of the *fabp* locus and the *EY02678* P-element insertion site. (D) Western blot of FABP and β-tubulin from adult fly head extracts of wild type or *fabp*^*EY02678*^ flies. (E) Western blot of Rh1 in flies from the head extracts of the indicated genotypes. *fabp wild type* (lane 1), *fabp*^*EY02678*^ mutants (lane 2), *fabp*^*EY02678*^ mutants rescued with *GMR-Gal4* driven *uas-fabp* expression (lane 3). Lanes 4 -6 show anti-Rh1 blots in the *ninaE*^*G69D*^*/+* background, with *fabp wild type* (lane 4), *fabp*^*EY0678*^ (lane 5), and *fabp*^*EY02678*^ mutants rescued with *GMR-Gal4/uas-fabp* (lane 6). (F) Quantification of relative Rh1 band intensities as normalized with β-tubulin. Error bars represent standard error (SE). Statistical significance was assessed through t-test. * = p<0.05, ** = p<0.005

To validate *fabp* RNAi results, we employed an *fabp* loss of function allele, *EY02678*, which has a P-element inserted near an exon-intron boundary (Figure 4C). This allele has strongly reduced FABP expression as assessed through western blot (Figure 4D). We found that *ninaE*^*G69D*^*/+* Rh1 levels increased in the *fabp*^*EY02678*^ *-/-* background (Figure 4E, F), validating the results with *fabp* RNAi. We further found that the loss of *fabp* increased Rh1 levels even in the *ninaE wild type* flies (Figure 4E, F). When we re-introduced *fabp* expression in *fabp* mutant flies using the eye specific *GMR-Gal4* driver, Rh1 protein levels were restored to those levels of wild type controls (Figures 4E, F). These results indicate that *fabp* affects general Rh1 protein levels.

### Regulation of Rh1 by *fabp* is light dependent

Since wild type Rh1 proteins are most notably degraded through light-dependent endocytosis (18-21), we examined whether *fabp* regulation of Rh1 was light-dependent. We found that *fabp* mutants showed higher Rh1 levels when the flies were reared under light. Such effect was not seen in flies that were reared in dark (Figure 5A, B).

**Figure 5:**
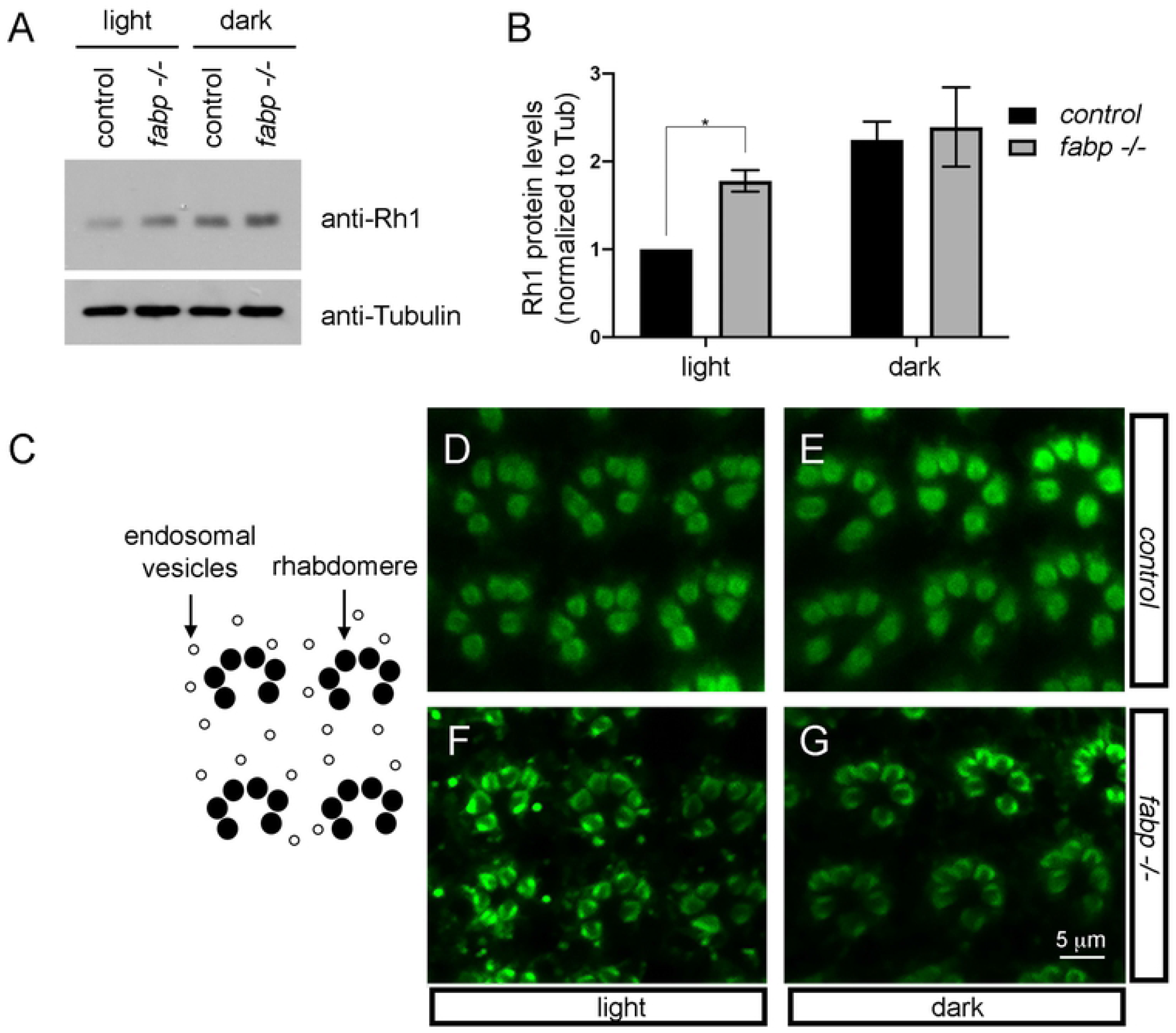
Rh1 regulation by *fabp* is light-dependent. (A) An anti-Rh1 western blot of control *fabp wild type* (lanes 1, 3) or *fabp*^*EY02678*^ *-/-* (lanes 2, 4) fly head extracts. The flies were either reared under constant light (lanes 1, 2) or in constant darkness (lanes 3, 4). The lower band shows an anti-Tubulin blot as a control. (B) Quantification of anti-Rh1 blot intensities, normalized to anti-Tubulin blots. Statistical significance was assessed through t-test. * = p<0.05. (C) A schematic diagram of adult *Drosophila* ommatidia, with rhabdomeres labeled as black circles, and endosomal vesicles as white circles. (D-G) Anti-Rh1 labeling of adult *Drosophila* ommatidia. (D, E) Control *fabp wild type* eyes. (F, G) *fabp*^*EY02678*^ *-/-* eyes. Fly samples (D, F) were reared under constant light before being processed for fixation and immuno-labeling. By contrast, samples (E, G) were reared under constant darkness before processing. The scale bar in G applies for images D – G.

To examine the pattern of Rh1 distribution in photoreceptors, we performed anti-Rh1 immuno-labeling in the adult *Drosophila* retina. In control wild type flies, Rh1 is predominantly detected in the rhabdomeres of R1 to R6 photoreceptor cells organized in a trapezoidal pattern (Figure 5C-E). In *fabp -/-* flies reared under light, however, there were additional anti-Rh1 signals in intracellular vesicles (Figure 5F).

Vesicular Rh1 signals reportedly appear in flies exposed to light, becoming even more prominent in mutants that have defects in Rh1 trafficking to the lysosome (24, 25). We found that vesicular Rh1 patterns in *fabp* -/- eyes were also light-dependent, as extra-rhabdomeric anti-Rh1 signals mostly disappeared in flies raised under constant darkness (Figure 5G). Together, these results suggest that light-activated Rh1 localize to intracellular vesicles, and these proteins are stabilized in *fabp* mutants.

### *fabp* mutants show light-dependent retinal degeneration that is suppressed in the *ninaE*^*G69D*^*/+* background

To examine whether *fabp* mutants affect retinal degeneration, we used Rh1-GFP flies with their photoreceptors labeled with green fluorescence. An intact ommatidia has R1-R6 photoreceptors arranged in a trapezoidal pattern that is readily visible as pseudopupils in live flies under low power microscopes (Figure 6A-D). Under standard conditions in which the flies were exposed to moderate light (see Methods), most wild type flies maintained this trapezoidal pattern of Rh1-GFP pseudopupils for the first thirty days after eclosion (Figure 6E, black line). *fabp*^*EY02678*^ *-/-* flies, on the other hand, showed signs of severe age-related retinal degeneration under otherwise identical conditions: Specifically, a few flies of this genotype began showing the loss of Rh1-GFP pseudopupils at day 15, with almost all examined flies having signs of retinal degeneration by day 28 (Figure 6E, red line; 6F). The difference between wild type controls and *fabp*^*EY02678*^ *-/-* was statistically significant (Log-rank test, p < 0.0001). Retinal degeneration in *fabp* mutants was light-dependent, as those reared in the dark did not exhibit signs of photoreceptor degeneration (Figure 6G). The light-dependent nature of photoreceptor degeneration correlated with *fabp*’s effect on Rh1 levels.

**Figure 6:**
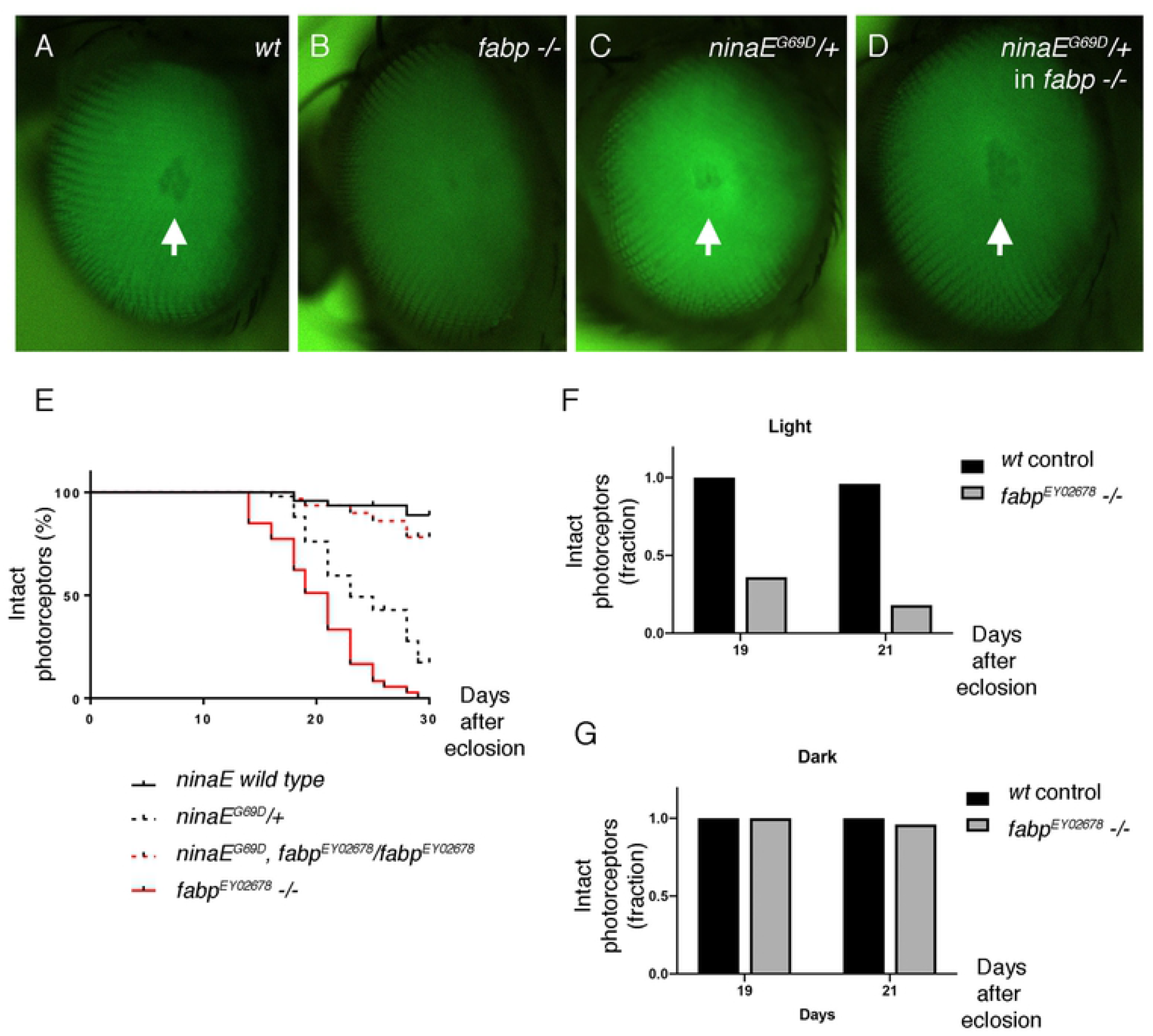
Retinal degeneration in *fabp* loss of function mutants. (A-D) Representative images of adult fly eyes of the indicated genotypes at 14 days after eclosion. White arrows point to the trapezoidal pattern of Rh1-GFP pseudopupils, which are indicative of intact photoreceptors. (E) The course of retinal degeneration in flies of the indicated genotypes, assessed through Rh1-GFP pseudopupils. Flies were reared under 1000 lux constant light. The y axis shows % of flies with intact photoreceptors. The x axis indicates the days after eclosion. *fabp*^*EY02678*^ mutant flies had accelerated retinal degeneration as compared to control *wild type* flies, with p < 0.0001 (Log-rank test). (F, G) Comparison of retinal degeneration state at 19 and 21 days after eclosion in flies reared under constant light (F), or under constant darkness (G). The y axis shows fraction of flies with intact photoreceptors. Black bars represent *fabp wild type* control flies, and the gray bars indicate *fabp*^*EY020866*^ -/- flies.

As reported previously, *ninaE*^*G69D*^*/+* flies showed age-related retinal degeneration that started occurring around day 17, with most flies exhibiting retinal degeneration at day 30 (Figure 6E, black dotted line). Surprisingly, flies containing *ninaE*^*G69D*^*/+* in the *fabp -/-* background had a significantly delayed course of retinal degeneration, with most flies still showing intact Rh1-GFP pseudopupils 30 days after eclosion. While surprising, such genetic interaction with *ninaE*^*G69D*^ is not unprecedented. Previous studies found that mutants that increase wild type Rh1 levels, such as *fatty acid transport protein* (*fatp*), cause severe retinal degeneration. Such retinal degeneration is suppressed in the *ninaE*^*G69D*^*/+* background (27).

## Discussion

The biological role of retinoic acid signaling is now well-delineated in vertebrates. However, the biological role and the mechanism of retinoid-mediated gene expression in *Drosophila* have remained unclear. Here, we showed that the expression of *Drosophila fabp* is regulated by vitamin A and retinoids. We found that *fabp* regulates Rh1 protein levels and loss of *fabp* results in retinal degeneration.

In *Drosophila*, the major phenotype associated with vitamin A deficiency is the loss of visual function. Significantly lower levels of Rh1 protein are detected under these conditions, as vitamin A deficiency would deplete the chromophore 11-cis 3-hydroxy retinal, which is normally required for proper Rh1 maturation. Our data presented here indicates that there is an additional layer of Rh1 regulation through retinoid-inducible *fabp*.

Several pieces of evidence presented here support the idea that *fabp* is involved in the endosomal/lysosomal degradation of light-activated Rh1. Specifically, we found that Rh1 levels increase in *fabp* mutants when flies were reared under light, but not when the flies were reared in constant darkness. Furthermore, immunohistochemical analysis shows that Rh1 accumulates in intracellular vesicles of *fabp* mutants only when the flies were reared under light. Since it is now well-documented that light-activated Rh1 undergoes endocytosis and lysosomal degradation (18-21), we interpret that *fabp* is specifically involved in this process.

We further note that the accelerated retinal degeneration phenotype of *fabp* mutants are reminiscent of other genetic conditions that result in endosomal accumulation of Rh1 in response to light. Examples of this type include mutations in *norpA, culd*, retromer complex proteins, and *fatty acid transport protein* (18, 25, 27, 45). As these genes normally regulate light-dependent internalization of Rh1, it is likely that excessive levels of light-activated Rh1 is contributing to the retinal degeneration phenotype under these conditions.

If it is indeed the excessive Rh1 levels in *fabp* mutants that are the cause of retinal degeneration, reduction of Rh1 in these flies would delay retinal degeneration. This is what we observed when we examined the *fabp* phenotype in the *ninaE*^*G69D*^*/+* background, which drastically reduces overall Rh1 levels. The way *fabp* and *ninaE*^*G69D*^ genetically interacted with each other was interesting in that *ninaE* ^*G69D*^*/+* flies normally show age-related retinal degeneration, which was also delayed in the *fabp* mutant background. This hints at the possibility that too low Rh1 levels is a contributing factor of retinal degeneration in *ninaE*^*G69D*^*/+* photoreceptors. Partial restoration functional Rh1 in the *fabp -/-* background appears to help delay retinal degeneration in *ninaE*^*G69D*^*/+* eyes.

In conclusion, we showed that *fabp* expression is regulated by retinoids and carotenoids in *Drosophila*. Our results indicate that retinoids not only serve as chromophores for rhodopsins, but have additional roles in regulating Rh1 protein levels. It remains to be examined whether mammalian CRAPBs similarly regulate rhodopsin levels and affect retinal degeneration in response to retinoic acids.

## Acknowledgements

We thank Drs. Vikke Weake, Jason Gerstner for fly lines and antibodies. This work was supported by the NIH grant R01 EY020866 to H.D.R.

## Author contributions

H.H. performed all experiments. H.H. and H.D.R. together designed experiments and analyzed data. H.D.R. wrote the manuscript draft with H.H.’s edits and inputs.

## Materials and Methods

### Fly Genetics

All fly crosses were maintained in 25 °C. Unless otherwise stated, flies were reared with a standard cornmeal-agar diet supplemented with molasses. Vitamin A deficient food was made by mixing 12 g yeast, 1.5 g agar, 7.5 g sucrose, 30 mg cholesterol, 3.75 ml of 1.15M Nippagin, 720 μl propionic acid in distilled water volume of 150 ml.

*Uas-fabp* had EGFP fused in frame with the *fabp’s* N-terminal coding sequence. *EGFP-fabp* was subcloned into the pUAST plasmid, and the resulting construct was injected by Best Gene, Inc., to generate the *uas-fabp* transgenic line.

We used the following flies that had been reported previously: *Rh1-Gal4* (49), *Rh1-GFP* (50), *ninaE*^*G69D*^ (9), *santa maria*^*1*^ (34), *ninaD*^*1*^ (31), *uas-dicer2* (51), *UAS-EGFP::Msp-300*^*KASH*^ (44). *fabp*^*CA06960*^ (52) and *fabp*^*EY02678*^ were obtained from the Bloomington *Drosophila* Stock Center (stock numbers #50808 and #15579, respectively).

The RNAi lines used are as follows: *uas-lacZ RNAi* (53), *uas-fabp RNAi* (Bloomington Stock Center # 34685), *uas-fatp RNAi* (Bloomington Stock Center # 55273), *uas-ninaB RNAi* (Bloomington Stock Center #34994), *uas-knrl RNAi* (Bloomington Stock Center # 36664), *uas-eg RNAi* (Bloomington # 35234). These lines were crossed to the female virgins of the genotype: *Rh1-Gal4; ninaE*^*G69D*^*/TM6B*. We collected non-TM6B progeny of these crosses to examine Rh1 protein and RNA.

### Photoreceptor-specific nuclear RNA extraction

We followed a published protocol to isolate Rh1-Gal4>UAS-EGFP::Msp-300^KASH^-positive nuclei (44). In brief, approximately 500 adult fly heads (from flies within 5 days of eclosion) per genotype were lysed in ice-cold nuclear isolation buffer (10 mM HEPES-KOH, pH 7.5; 2.5 mM MgCl_2_; 10 mM KCl) with a dounce homogenizer. The homogenate was filtered through a 40µm Flowmi cell strainer (WVR, cat. #BAH136800040), and the filtrate was incubated with anti-EGFP-coupled protein G Dynabeads (Invitrogen, cat. #10003D) for 1 hour at 4°C. The beads were collected using a magnetic microcentrifuge tube holder (Sigma, cat. #Z740155). Following washes with wash buffer (PBS, pH 7.4; 2.5mM MgCl_2_), the beads were resuspended in a final volume of 150µL of wash buffer. Then the post-isolation nuclei were suspended in 1mL of Trizol reagent (Life Technologies, cat. #15596018) for RNA extraction following standard procedures. Prior to RNA precipitation with isopropanol, 0.3M sodium acetate and glycogen were added to facilitate visualization of the RNA pellet. We then suspended the pellet in RNAse-free water and purified it using a Qiagen RNeasy MinElute cleanup kit (Qiagen, cat. #74204) following standard protocols.

### Preparation of cDNA libraries, RNA-seq and data processing

The NYU Genome Technology Center performed library preparation and RNA sequencing. We quantified RNA on an Agilent 2100 BioAnalyzer (Agilent, cat. #G2939BA). For cDNA library preparation and ribodepletion, we utilized a custom *Drosophila* Nugen Ovation Trio low-input library preparation kit (Tecan Genomics), using approximately 20 ng total RNA per sample. For sequencing, we performed paired-end 50bp sequencing of samples on an Illumina NovaSeq 6000 platform (Illumina, cat. #20012850) using half of a 100 cycle SP flow cell (Illumina, cat. #20027464). We used the bcl2fastq2 Conversion software (v2.20) to convert per-cycle BCL base call files outputted by the sequencing instrument (RTA v3.4.4) into the fastq format in order to generate per-read per-sample fastq files. For subsequent data processing steps, we used the Seq-N-Slide automated workflow developed by Igor Dolgalev (https://github.com/igordot/sns). For read mapping, we used the alignment program STAR (v2.6.1d) to map reads of each sample to the *Drosophila melanogaster* reference genome dm6, and for quality control we used the application Fastq Screen (v0.13.0) to check for contaminating sequences. We employed featureCounts (Subread package v1.6.3) to generate matrices of read counts for annotated genomic features. For differential gene statistical comparisons between groups of samples contrasted by genotype, we used the DESeq2 package (R v3.6.1) in the R statistical programming environment. We excluded genes with baseMean counts less than 300 so as to avoid artifacts due to varying extent of nuclei purification.

### Immunofluorescence and Western Blots

We followed standard protocols for western blots and whole mount immuno-labeling experiments using the following primary antibodies: Mouse monoclonal 4C5 anti-Rh1 (Developmental Studies Hybridoma Bank, used at 1:5000 for western blots), anti-β tubulin antibody (Covance #MMS-410P), Rabbit anti-GFP (Invitrogen #A-6455), anti-FABP antibody (54).

### RT-PCR

We performed qRT-PCR using Power SYBR green master mix kit (Thermo Fisher). The primer sequences are as follows:

Rpl15F: AGGATGCACTTATGGCAAGC

Rpl15R: GCGCAATCCAATACGAGTTC

FatpF: CTCCCGGTGAGTGCAATAGCTT

FatpR: GCGGTGTGGTACAAAGGCAA

Arr1F: CATGAACAGGCGTGATTTTGTAG

Arr1R: TTCTGGCGCACGTACTCATC

Arr2F: TCGATGGAGTGATTGTGGTGG

Arr2R: GCGACCATAGCGATAGGTGG

Fabp-1F: CCGAGGTCTCAGTGTGCTC

Fabp-1R: CCGAGGTCTCAGTGTGCTC

Fabp-2F: CACAGTGGAGGTGACCTTGG

Fabp-2R: GATGCTCTTGACGTTGCGAC

TubF: CTCAGTGCTCGATGTTGTCC

TubR: GCCAAGGGAGTGTGTGAGTT

### Retinal degeneration assay

We performed all retinal degeneration assays in the *cn, bw -/-* background to eliminate eye pigments that otherwise affect the course of retinal degeneration. The flies were incubated in the 25 °C incubator with 1000 lux of light. For retinal degeneration assays under constant darkness, the flies were reared in an enclosed cardboard box in the 25 °C incubator. Retinal degeneration was assessed based on green fluorescent pseudopupils originating from the *Rh1-GFP* transgene. We interpreted clear trapezoidal pseudopupils as evidence in intact photoreceptors, while its disappearance was construed as a sign of retinal degeneration. The number of flies analyzed for each genotype in Figure 6E is as follows:

*wild type*, 48 flies; *fabp*^*EY02678*^, 52 flies; *ninaE*^*G69D*^*/+*, 50 flies; *ninaE*^*G69D*^, *fabp*^*EY02678*^*/ fabp*^*EY02678*^, 32 flies.

For Figures 6F and G, 50 flies were analyzed for each genotype.

### Quantification and statistics

To quantify proteins in gels, we measured average pixel intensities of western blot bands using Image J, and normalized them to anti-β tubulin bands. Graphs were generated after at least three independent measurements and p values were calculated using a paired t-test. For retinal degeneration assays, we used the Log-rank (Mantel-Cox) text. Graphs were made using the *Graphpad Prism* program. All error bars represent SEM (Standard error of the mean).

## References

1. Yau KW, Hardie, R.C. Phototransduction motifs and variations. Cell. 2009;139(2):246–64.

2. O’Tousa JE, Baehr, W., Martin, R.L., Hirsh, J., Pak, W.L., Applebury, M.L. The Drosophila ninaE gene encodes an opsin. Cell. 1985;40(4):877–82.

3. Zuker CS, Cowman, A.F., Rubin, G.M. Isolation and structure of a rhodopsin gene from D. melanogaster. Cell. 1985;40(4):851–8.

4. Montell C. Drosophila visual transduction. Trends Neurosci. 2012;35(6):356–63.

5. Nichols R, Pak WL. Characterization of Drosophila melanogaster rhodopsin. J Biol Chem. 1985;260(23):12670–4.

6. Seki T, Isono K, Ito M, Katsuta Y. Flies in the group Cyclorrhapha use (3S)-3-hydroxyretinal as a unique visual pigment chromophore. Eur J Biochem. 1994;226(2):691–6.

7. Dewett D, Lam-Kamath K, Poupault C, Khurana H, Rister J. Mechanisms of vitamin A metabolism and deficiency in the mammalian and fly visual system. Dev Biol. 2021.

8. Scavarda NJ, O’Tousa, J., Pak, W.L. Drosophila locus with gene-dosage effects on rhodopsin. Proc Natl Acad Sci USA. 1983;80(14):4441–5.

9. Colley NJ, Cassill, J.A., Baker, E.K., Zuker, C.S. Defective intracellular transport is the molecular basis of rhodopsin-dependent dominant retinal degeneration. Proc Natl Acad Sci USA. 1995;92(7):3070–4.

10. Kurada P, O’Tousa, J.E. Retinal degeneration caused by dominant rhodopsin mutations in Drosophila. Neuron. 1995;14(3):571–9.

11. Dryja T, McGee, T.L., Reichel, E., Hahn, L.B., Cowley, G.S., Yandell, D.W., Sandberg, M.A., Berson, E.L. A point mutation of the rhodopsin gene in one form of retinitis pigmentosa. Nature. 1990;343(6256):364–6.

12. Sung CH, Davenport, C.M., Hennessey, J.C., Maumenee, I.H., Jacobson, S.G., Heckenlively, J.R., Nowakowski, R., Fishman, G., Gouras, P., Nathans, J. Rhdopsin mutations in autosomal dominant retinitis pigmentosa. Proc Natl Acad Sci USA. 1991;88(15):6481–5.

13. Ryoo HD, Domingos PM, Kang MJ, Steller H. Unfolded protein response in a Drosophila model for retinal degeneration. Embo j. 2007;26(1):242–52.

14. Kang M-J, Ryoo, H.D. Suppression of retinal degeneration in Drosophila by stimulation of ER-Associated Degradation. Proc Natl Acad Sci USA. 2009;106(40):17043–8.

15. Lin JH, Li, H., Yasumura, D., Cohen, H.R., Zhang, C., Panning, B., Shokat, K.M., Lavail, M.M., Walter, P. IRE1 signaling affects cell fate during the unfolded protein response. Science. 2007;318(5852):944–9.

16. Xu J, Zhao H, Wang T. Suppression of retinal degeneration by two novel ERAD ubiquitin E3 ligases SORDD1/2 in Drosophila. PLoS Genet. 2020;16(11):e1009172.

17. Dolph PJ, Ranganathan R, Colley NJ, Hardy RW, Socolich M, Zuker CS. Arrestin function in inactivation of G protein-coupled receptor rhodopsin in vivo. Science. 1993;260(5116):1910–6.

18. Alloway PG, Howard L, Dolph PJ. The formation of stable rhodopsin-arrestin complexes induces apoptosis and photoreceptor cell degeneration. Neuron. 2000;28(1):129–38.

19. Orem NR, Dolph, P.J. Epitope masking of rhabdomeric rhodopsin during endocytosis-induced retinal degeneration. Mol Vis. 2002;8:455–61.

20. Satoh AK, Ready DF. Arrestin1 mediates light-dependent rhodopsin endocytosis and cell survival. Curr Biol. 2005;15(19):1722–33.

21. Satoh AK, Xia H, Yan L, Liu CH, Hardie RC, Ready DF. Arrestin translocation is stoichiometric to rhodopsin isomerization and accelerated by phototransduction in Drosophila photoreceptors. Neuron. 2010;67(6):997–1008.

22. Xu H, Lee SJ, Suzuki E, Dugan KD, Stoddard A, Li HS, et al. A lysosomal tetraspanin associated with retinal degeneration identified via a genome-wide screen. Embo j. 2004;23(4):811–22.

23. Orem NR, Xia L, Dolph PJ. An essential role for endocytosis of rhodopsin through interaction of visual arrestin with the AP-2 adaptor. J Cell Sci. 2006;119(Pt 15):3141–8.

24. Chincore Y, Mitra, A., Dolph, P.J. Accumulation of rhodopsin in late endosomes triggers photoreceptor cell degeneration. PLoS Genet. 2009;5(2):e1000377.

25. Wang S, Tan, K.L., Agosto, M.A., Xiong, B., Yamamoto, S., Sandoval, H., Jaiswal, M., Bayat, V., Zhang, K., Charng, W.L., David, G., Duraine, L., Venkatachalam, K., Wensel, T.G., Bellen, H.J. The retromer complex is required for rhodopsin recycling and its loss leads to photoreceptor degeneration. PLoS Biology. 2014;12(4):e1001847.

26. Hebbar S, Lehmann M, Behrens S, Hälsig C, Leng W, Yuan M, et al. Mutations in the splicing regulator Prp31 lead to retinal degeneration in Drosophila. Biol Open. 2021;10(1).

27. Dourlen P, Bertin B, Chatelain G, Robin M, Napoletano F, Roux MJ, et al. Drosophila fatty acid transport protein regulates rhodopsin-1 metabolism and is required for photoreceptor neuron survival. PLoS Genet. 2012;8(7):e1002833.

28. Chuang JZ, Vega C, Jun W, Sung CH. Structural and functional impairment of endocytic pathways by retinitis pigmentosa mutant rhodopsin-arrestin complexes. J Clin Invest. 2004;114(1):131–40.

29. Chen J, Shi G, Concepcion FA, Xie G, Oprian D, Chen J. Stable rhodopsin/arrestin complex leads to retinal degeneration in a transgenic mouse model of autosomal dominant retinitis pigmentosa. J Neurosci. 2006;26(46):11929–37.

30. Harris WA, Ready, D.F., Lipson, E.D., Hudspeth, A.J., Stark, W.S. Vitamin A deprivation and Drosophila photopigments. Nature. 1977;266(5603):648–50.

31. Gu G, Yang, J., Mitchell, K.A., O’Tousa, J.E. Drosophila ninaB and ninaD act outside of retina to produce rhodopsin chromophore. J Biol Chem. 2004;279(18):18608–13.

32. Wang T, Montell, C. Rhodopsin formation in Drosophila is dependent on the PINTA retinoid-binding protein. J Neurosci. 2005;25(21):5187–94.

33. Ahmad ST, Joyce, M.V., Boggess, B., O’Tousa, J.E. The role of Drosophila ninaG oxidoreductase in visual pigment chromophore biogenesis. J Biol Chem. 2006;281(14):9205–9.

34. Wang T, Jiao, Y., Montell, C. Dissection of the pathway required for generation of vitamin A and for Drosophila phototransduction. J Cell Biol. 2007;177(2):305–16.

35. Mangelsdorf DJ, Evans, R.M. The RXR heterodimers and orphan receptors. Cell. 1995;83(6):841–50.

36. Kleywegt GJ, Bergfors, T., Senn, H., Le Motte, P., Gsell, B., Shudo, K., Jones, T.A. Crystal structures of cellular retinoic acid binding proteins I and II in complex with all-trans-retinoic acid and a synthetic retinoid. Structure. 1994;2(12):1241–58.

37. Napoli JL. Cellular retinoid binding-proteins, CRBP, CRABP, FABP5: Effects on retinoid metabolism, function and related diseases. Pharmacol Ther. 2017;173:19–33.

38. Durand B, Saunders, M., Leroy, P., Leid, M., Chambon, P. All-trans and 9-cis retinoic acid induction of CRABPII transcription is mediated by RAR-RXR heterodimers bound to DR1 and DR2 repeated motifs. Cell. 1992;71(1):73–85.

39. King-Jones K, Thummel, C.S. Nuclear receptors-a perspective from Drosophila. Nat Rev Genet. 2005;6(4):311–23.

40. Picking WL, Chen, D.M., Lee, R.D., Vogt, M.E., Polizzi, J.L., Marietta, R.G., Stark, W.S. Control of Drosophila opsin gene expression by carotenoids and retinoic acid: northern and western analysis. Exp Eye Res. 1996;63(5):493–500.

41. Shim K, Picking, W.L, Kutty, R.K., Thomas, C.F., Wiggert, B.N., Stark, W.S. Control of Drosophila retinoid and fatty acid binding glycoprotein expression by retinoids and retinoic acid: northern, western and immunocytochemical analysis. Exp Eye Res. 1997;65(5):717–27.

42. Huang HW, Brown, B., Chung, J., Domingos, P.M., Ryoo, H.D. highroad is a carboxypeptidase induced by retinoids to clear mutant Rhodopsin-1 in Drosophila Retinitis Pigmentosa models. Cell Rep. 2018;22(6):1384–91.

43. Hall H, Medina P, Cooper DA, Escobedo SE, Rounds J, Brennan KJ, et al. Transcriptome profiling of aging Drosophila photoreceptors reveals gene expression trends that correlate with visual senescence. BMC Genomics. 2017;18(1):894.

44. Ma J, Weake VM. Affinity-based isolation of tagged nuclei from Drosophila tissues for gene expression analysis. J Vis Exp. 2014(85).

45. Xu Y, Wang T. CULD is required for rhodopsin and TRPL channel endocytic trafficking and survival of photoreceptor cells. J Cell Sci. 2016;129(2):394–405.

46. Giguere V, Lyn, S., Yip, P., Siu, C.H., Amin, S. Molecular cloning of cDNA encoding a second cellular retinoic acid-binding protein. Proc Natl Acad Sci USA. 1990;87(16):6233–7.

47. Shaw N, Elholm, M., Noy, N. Retinoic acid is a high affinity selective ligand for the peroxisome proliferator-activated receptor beta/delta. J Biol Chem. 2003;278(43):41589–92.

48. von Lintig J, Dreher, A., Kiefer, C., Wernet, M.F., Vogt, K. Analysis of the blind Drosophila mutant ninaB identifies the gene encoding the key enzyme for vitamin A formation invivo. Proc Natl Acad Sci USA. 2001;98(3):1130–5.

49. Mollereau B, Wernet MF, Beaufils P, Killian D, Pichaud F, Kuhnlein R, et al. A green fluorescent protein enhancer trap screen in Drosophila photoreceptor cells. Mech Dev. 2000;93(1-2):151–60.

50. Pichaud F, Desplan C. A new visualization approach for identifying mutations that affect differentiation and organization of the Drosophila ommatidia. Development. 2001;128(6):815–26.

51. Dietzl G, Chen, D., Schnorrer, F., Su, K.C., Barinova, Y., Fellner, M., Gaser, B., Kinsey, K., Oppel, S., Scheiblauer, S., Couto, A., Marra, V., Keleman, K., Dickson, B.J. A genome-wide transgenic RNAi library for conditional gene inactivation in Drosophila. Nature. 2007;448(7150):151–6.

52. Buszczak M, Paterno, S., Lighthouse, D., Bachman, J., Planck, J., Owen, S., Skora, A.D., Nystul, T.G., Ohlstein, B., Allen, A., Wilhelm, J.E., Murphy, T.D., Levis, R.W., Matunis, E., Srivali, N., Hoskins, R.A., Spradling, A.C. The Carnegie protein trap library: A versatile tool for Drosophila development studies. Genetics. 2007;175(3):1505–31.

53. Kennerdell JR, Carthew RW. Heritable gene silencing in Drosophila using double-stranded RNA. Nat Biotechnol. 2000;18(8):896–8.

54. Gerstner JR, Vanderheyden WM, Shaw PJ, Landry CF, Yin JC. Fatty-acid binding proteins modulate sleep and enhance long-term memory consolidation in Drosophila. PLoS One. 2011;6(1):e15890.

